# Zebrafish *follistatin-like 1b* regulates cardiac contraction during early development

**DOI:** 10.1101/2020.06.23.167957

**Authors:** Xin-Xin I. Zeng, Karen Ocorr, Erik J. Ensberg, P. Duc si Dong

## Abstract

Extracellular glycoprotein Follistatin-like protein 1 (FSTL1) has been reported to be involved in multiple signaling pathways and biological processes during development and disease. In addition, some mouse studies have suggested an inhibitory role of FSTL1 in BMP signaling, in certain context. Yet, whether FSTL1 has biological functions in early heart development are largely unknown. In our functional studies, CRISPR/Cas9 genome editing was used to create *fstl1a* and *fstl1b* insertion/deletion mutations in zebrafish and our results suggest that *fstl1b* is important in regulating heart development during early embryogenesis. *in situ* hybridization revealed that *fstl1b* transcripts are expressed in the developing zebrafish heart, and *fstl1b* homozygous (-/-) mutant exhibited pericardial edema and showed significantly reduced contractility in the ventricle, with incomplete penetrance. We found that zebrafish *fstl1b (-/-)* mutant hearts formed a collapsed ventricle with strong reductions in the sarcomere structure protein alpha-actinin, although the number of cardiomyocytes was comparable to wild type control siblings. Further, normal levels of *fstl1b* seems to prevent the expansion of Bmp signaling into ventricular myocytes, which is consistent with the previously established model of Bmp-dependent cardiac contraction. Together these results reveal the zebrafish ortholog *fstl1b* regulates sarcomere structure and cardiac contractile function. In vertebrate developing heart, it might function as a key component in a novel pathway that constrains Bmp signaling from ventricular myocytes.

## Introduction

Follistatin-like protein 1 (FSTL1) is an extracellular glycoprotein belonging to the secreted protein acid and rich in cysteine (SPARC) family containing both extracellular calcium-binding and follistatin-like domains (1, 2). FSTL1 is involved in multiple signaling pathways and biological processes during development and disease, including cardiovascular disease, cancer, and arthritis (3, 4). Functional disruption of Fstl1 in mice has demonstrated its important roles in early development and organogenesis, such as lung (5, 6), skeletal tissues (6), and ureter (7). These studies also suggested an inhibitory role of FSTL1 in BMP signaling (5, 6). Further, FSTL1 protein seems to have a cardioprotective role. Although the mechanism of action remains elusive, FSTL1 attenuated hypertrophy following pressure overload (8), and prevented myocardial ischemia/reperfusion injury in a mouse or pig model of ischemia/reperfusion (9). Most recently, FSTL1 administered therapeutically has been shown to have a pronounced ability to stimulate regeneration following myocardial infarction. Treating experimental animals (mouse and pig) with FSTL1 after myocardial infarction progressively restored heart function, at least in part by stimulating replication of normally non-dividing heart muscle cells (10). Despite these studies, its role in regulating cardiac development has been less explored, probably due to homozygous lethality in the mouse model (5, 6, 11).

In the field of cardiovascular research, due to a unique combination of visual access to the early embryonic cardiovascular system and established tools and methods for genetic analysis, the zebrafish system is recognized as an ideal model organism for investigating the molecular mechanisms underlying heart and blood vessel formation (12-14). The zebrafish embryo does not require a functional cardiovascular system for its survival through 5 days post fertilization (dpf) (15, 16), thus cardiovascular defects can be analyzed in detail throughout embryogenesis. Due to genome duplication, the zebrafish genome contains two orthologs of the human *FSTL1* gene, *fstl1a* and *fstl1b* (17). In many instances, zebrafish orthologs have subtly diverged in structure and function (18, 19). Since the introduction of new genome editing tools such as ZFNs (zinc finger nucleases), TALENs (transcription activator-like effector nucleases), and more recently, the CRISPR (clustered regularly interspaced short palindromic repeats)/CRISPR-associated (Cas) system has greatly expanded the ability to knock out genes in various animal models with high efficiency (20-22), including zebrafish (23-26). Here, we used the CRISPR/Cas9 system as a genome editing tool and efficient protocols that have been generated for zebrafish (24, 26, 27), to successfully generate deletion and insertion mutations in the *fstl1a* and *fstl1b* loci.

We used these zebrafish *fstl1a* and *fstl1b* mutants to explore the regulatory relationship between Fstl and Bmp signaling in vertebrate heart development. Here we show that *fstl1b* homozygous mutants exhibited substantial reductions of cardiac ventricular contraction by quantitative measurements. Consistently, sarcomere assembly was destabilized in the mutant cardiomyocytes by immunofluorescence staining analysis. Furthermore, *bmp4* and phosphorylated-Smad expression seems to be elevated in *fstl1b* homozygous mutant hearts. Thus, the roles of Fstl1 and BMP signaling interaction might reveal a novel mechanism for the regulation of the integrity of cardiac muscle structure and cardiac cell function.

## Results

### Zebrafish *fstl1a* and *fstl1b* CRISPR/Cas9 generated mutants show relatively normal embryonic development

We performed targeted mutagenesis using CRISPR/Cas9 genome editing (24, 26, 27), to induce indel mutations in both *fstl1a* and *fstl1b* genes with potential loss-of-function effects. We designed two single-stranded gRNAs (sgRNAs) composed of the targeting sequence followed by the PAM sequence that guides the Cas9 nuclease to the desired genomic locus of zebrafish *fstl1a* and *fstl1b* (Fig. 1A, D). The sgRNAs were designed to target the *fstl1a* gene’s Exon 2 region, and *fstl1b* gene’s Exon 4 region (Fig. 1D; Supplementary Table 1). The sgRNAs were obtained by in *vitro* transcription directly from synthesized oligonucleotides. Subsequently, the sgRNA was coinjected into fertilized one-cell stage embryos along with Cas9 protein from *Streptococcus pyogenes* (EnGen^®^ Spy Cas9 NLS, NEB #M0646), in which two nuclear localization signals (NLSs) are added to target it to the nucleus (28) (Fig. 1B). Injected embryos were analyzed at 24 hours post-fertilization (hpf) for genome modifications of the target locus. PCR products including the target site were amplified and analyzed using high concentration (>4%) agarose gel electrophoresis as an efficient technology for indel detection and to distinguish the size of PCR products between the heterozygous mutant and the wild-type genotypes (not shown). Sanger sequence analysis confirmed that the loss of the respective restriction site was due to mutations at the target sites in the sgRNA/Cas9-injected embryos (Fig. 1D). Further, we have created several lines with unique mutations in *fstl1a* and *fstl1b*, and these lines were outcrossed and raised to the F8 generation and beyond prior to use, which significantly reduces the incidence of potential off-target effects (Fig. 1C; Supplemental Table 1).

**Figure 1.**
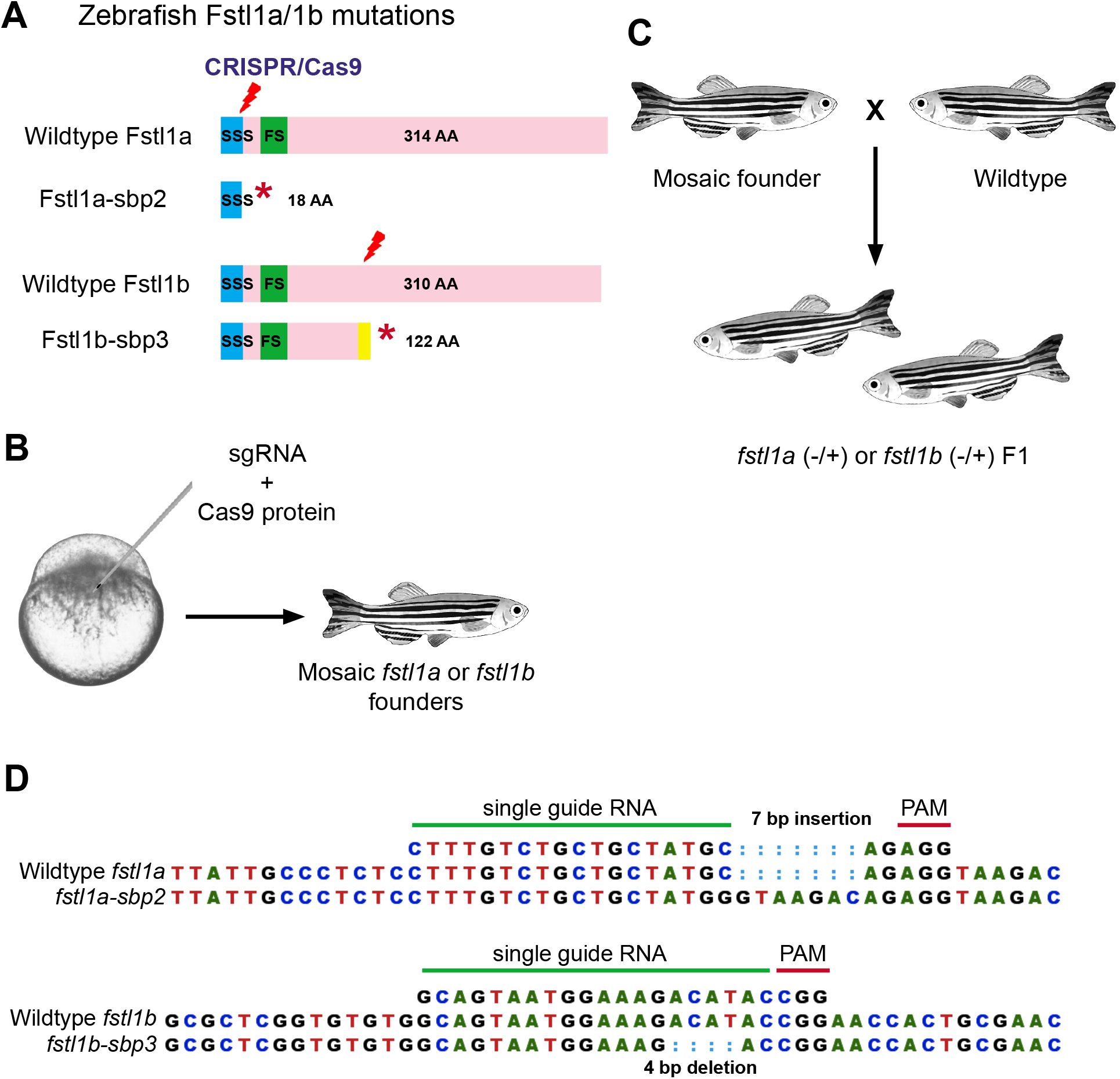
CRISPR/Cas9 introduce mutations in zebrafish *fstl1a* and *fstl1b* genes. (A) Schematic of the zebrafish Fstl1a and Fstl1b locus and CRISPR/Cas9 targeted region (red arrowheads). * denote stop codon. (B and C) Injection and breeding paradigm to generate Fstl1a and Fstl1b heterozygosity. (D) Top panel, Sanger sequencing of *fstl1a-sbp2* mutation allele showing 7 base pair insertion with one single nucleotide change (C>G) at the 5’ ends as well; Bottom panel, Sanger sequencing of *fstl1b-sbp3* mutation allele showing 4 base pair deletion. The green line indicates the sequences of each single guide RNA, and the red line highlights the corresponding PAM region.

We began our functional and phenotypical analysis of the mutant alleles *fstl1a-sbp2* and *fstl1b-sbp3* (Fig. 1D; Supplemental Table 1). Zebrafish *fstl1a* heterozygous or homozygous mutants (generated by *fstl1a* heterozygous parents incross) appear free of developmental and cardiac abnormalities (n=63; Fig. 2A). Zebrafish *fstl1b* heterozygous mutants (generated by *fstl1b* heterozygous parents incross) also develop normally through early development (n=30), In contrast, approximately half (37%, n=24) of *fstl1b* homozygous embryos (generated by *fstl1b* heterozygous parents) exhibited mild cardiac edema starting at 60 hpf (Fig. 2B, lower panel in the homozygous category, arrow shows epicardial edema), indicating that this particular *fstl1b* allele might exhibit incomplete penetrance. We have confirmed that both single *fstl1a* and *fstl1b* homozygous (generated by heterozygous parents) mutants are viable and survive to adulthood, *fstl1a* homozygous has one quarter adult fish in ratio being identified at 4-month-old (20 homozygous in total 81 adult fish); however, only approximal 10% *fstl1b* homozygous adult fish survived (12 homozygous in total 123 adult fish).

**Figure 2.**
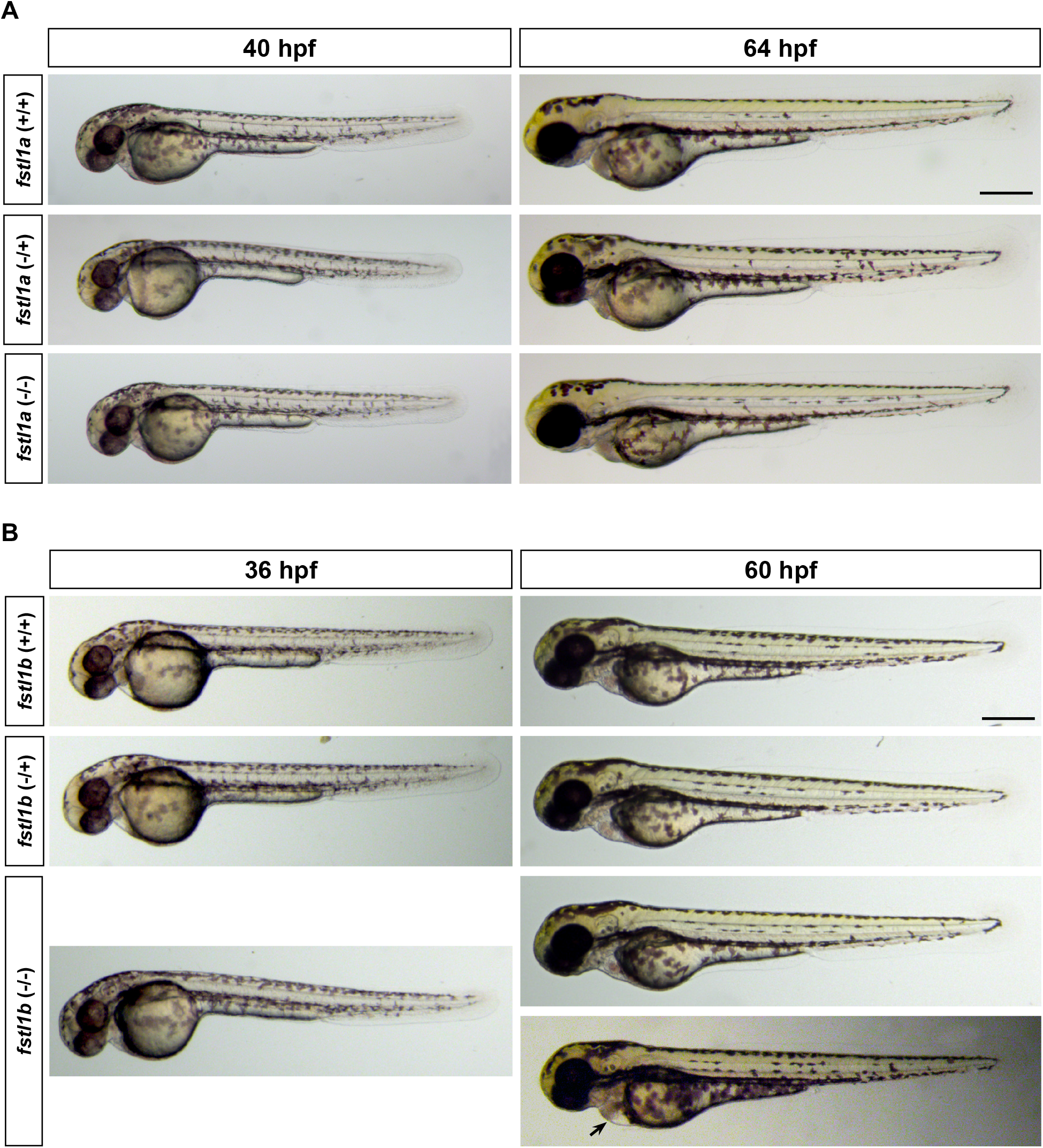
Zebrafish *fstl1a* and *fstl1b* mutants exhibit relatively normal embryonic development. Phenotypical analysis of mutant fish alleles *fstl1a-sbp2* and *fstl1b-sbp3.* (A) Representative bright-field (40; 64 hpf) images of wild-type, *fstl1a* heterozygous, and homozygous zebrafish embryos. Lateral view, head to the left. Zebrafish *fstl1a* homozygous mutant develop normally through early development. (B) Representative bright-field (36; 60 hpf) images of wildtype, *fstl1b* heterozygous, and homozygous zebrafish embryos. Lateral view, head to the left. Zebrafish *fstl1b* homozygous mutant develop normally through early development, and less than half (37%, n=24) of homozygous embryos start to exhibit mild cardiac edema at 60 hpf (arrow). hpf, hours post fertilization. Scale bars: 300 μm.

### Zebrafish *fstl1b* is expressed in the ventricular region during development and is important for cardiac contractility

The cardiac edema phenotypes observed in *fstl1b* (-/-) homozygous mutant (*sbp3* allele) embryos suggested a potential abnormal heart development and cardiac dysfunction (Fig. 2B, arrow). We have previously developed a zebrafish heart preparation and movement analysis algorithm that allows the quantification of functional parameters (29, 30). This methodology combines highspeed optical recording of beating hearts with a robust, semi-automated analysis SOHA (Semiautomated Optical Heartbeat Analysis) to accurately detect and quantify, on a beat-to-beat basis, heart period, systolic and diastolic chamber size, fractional area change (FAC) and cardiac arrhythmicity. We performed a detailed analysis of hearts from 3-day-old zebrafish larva, aimed to detect potentially subtle but distinct cardiac deficits in *fstl1a* and *fstl1b* mutant zebrafish. Compared to wild-type siblings (Fig. 3A), *fstl1b* homozygous embryos (*fstl1b-sbp3* allele was analyzed in this set of experiments), which have pericardial edema, exhibit a much more linear heart tube at 72 hpf (Fig. 3B, Supplemental Movie 1), at which stage both chambers should have completed the looping process during normal development (15, 31, 32) (Fig. 3A). Strikingly, the cardiac contractility, measured as fractional area change (End Diastolic Area - End Systolic Area/ End Diastolic Area), was significantly reduced in ventricles from *fstl1b* homozygous mutants due to this diastolic dysfunction (Fig. 3C). This reduction in contractility was associated with a reduction in overall size of the ventricles during diastole suggesting diastolic dysfunction. Contractility was also reduced in atria from *fstl1b* homozygous mutants, despite the fact that the overall size of the atria did not change significantly (Fig. 3D). In parallel, no obvious cardiac dysfunction was detected in *fstl1a* homozygous mutants (*fstl1a-sbp2* allele was analyzed in this set of experiments, Supplemental Fig. 1).

**Figure 3.**
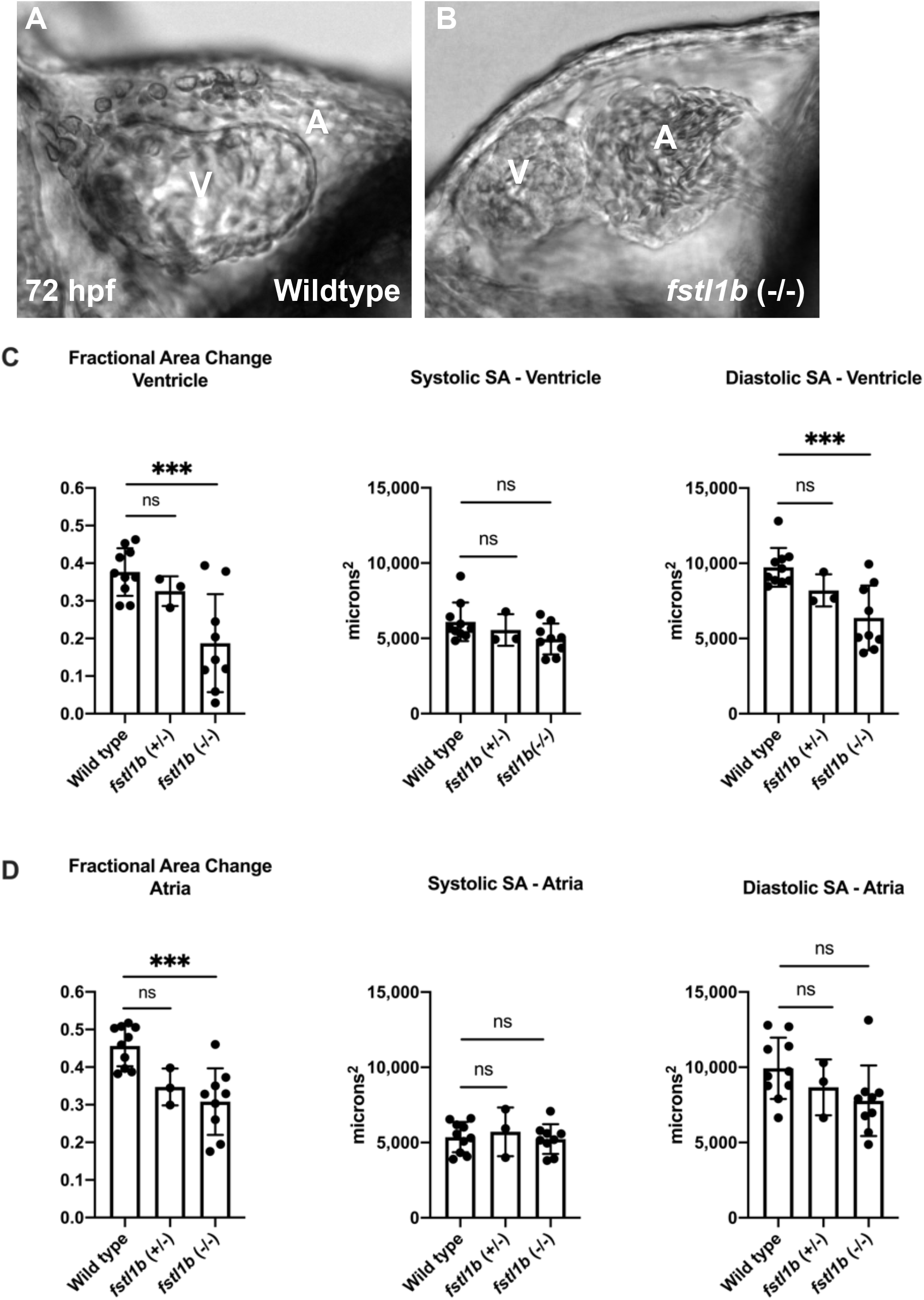
Loss of *fstl1b* function causes cardiac disfunction, particularly leads to strongly reduced contractility in the ventricle. At 72 hours post fertilization (hpf), representative still images from high speed movies showing zebrafish hearts. Anterior is to the left and dorsal is on the bottom in all images. (A) Wildtype embryo *fstl1b* (+/+) exhibit a well-developed ventricle (V) superimposed over the atrium (A) as a result of normal looping. (B) Homozygous mutant embryos *fstl1b* (-/-) exhibit a linear heart tube that failed to complete looping, as well as hypertrophic and restricted ventricles associated with edema of the pericardial sac (V, Ventricle; A, Atrium). We quantified cardiac function in *fstl1b* mutants using high-speed imaging and SOHA (Semiautomated Optical Heartbeat Analysis). Strikingly, we found that, the cardiac contractility measured as fractional area change (End Diastolic Area - End Systolic Area/ End Diastolic Area), was significantly reduced in ventricles from *fstl1b* homozygous mutants due to significantly altered diastolic dysfunction (C), and atrial parameters were less affected (D). All hearts were imaged at room temperature (20-21°C) in a double-blind manner; genotypes were identified after filming and movie analyses had been completed. Statistical analyses were performed using Prism software (GraphPad). Significance was determined using one-way ANOVA and Dunnett’s multiple comparisons post hoc test. (ns, not significant; ***, p<0.0001).

To examine the expression patterns of *fstl1a* and *fstl1b* during heart development, we took advantage of a previously established fluorescent *in situ* hybridization protocol to sensitively detect the endogenous transcripts and reveal both genes’ expression domains in high-resolution. Further, we also used MF20 staining to landmark the location of mature cardiomyocytes. At 48 hpf, we found that *fstl1b* is broadly expressed in the ventricle, the atrium, and the cardiac outflow tract regions (Fig. 4D-F; D’-F’; D’’-F’’). Interestingly, there was no detectable level of *fstl1a* expression in the heart or around the cardiac tissues during the similar developmental stages. (48 hpf; not shown). Taken together, these results indicate that zebrafish ortholog *fstl1a* and *fstl1b* may have distinct roles during early embryonic development: *fstl1b* might be the relevant one that play an essential role in regulating the ventricular muscle development and function.

**Figure 4.**
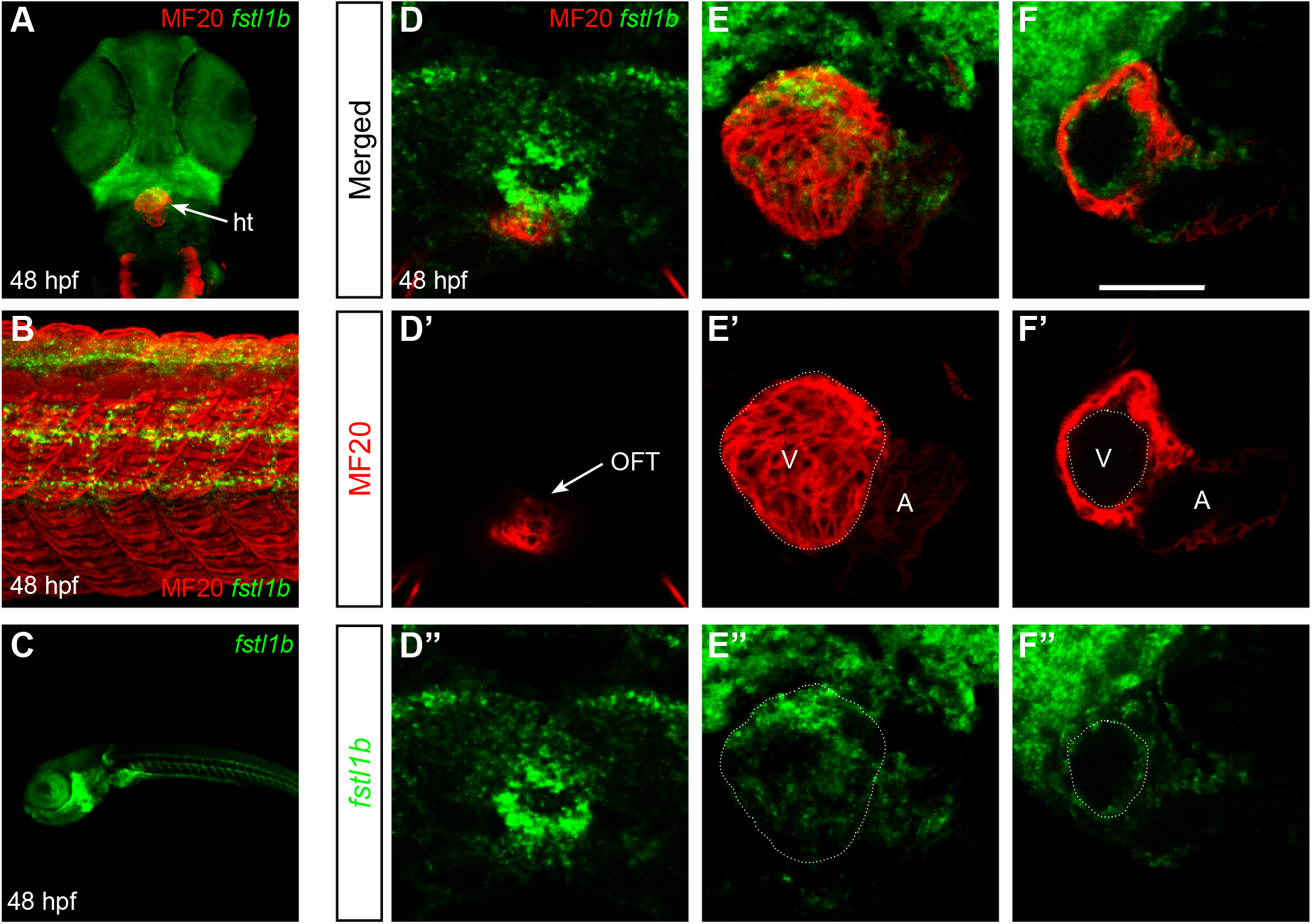
Whole-mount fluorescent *in situ* hybridization demonstrates the expression patterns of *fstl1b* in the zebrafish embryonic heart. (A,B,C) At 48 hpf, confocal rendered 3D images show expression of *fstl1b* thought the entire body (C, lateral view, head is the left), and demonstrated its colocalization in the heart (A, ventral view, head is to top), the truck region (lateral view, anterior is to the left) when co-stained with MF20 antibody (green, *fstl1b* antisense probe; red, MF20 antibody). ht, heart. (D-F, merged view; D’-F’, MF20 channel; D”-F”, *fstl1b* fluorescent in situ channel’’) Single confocal slices through zebrafish hearts at 48 hpf, arterial pole up. Whole-mount fluorescent *in situ* hybridization shows that *fstl1b* (green) is expressed in the region where the pole of OFT cells are thought to reside (D’, arrow), as well as in the ventricular myocardium tissues (dotted circle outlines the ventricle). Differentiated myocardium is marked by MF20 (red). (V, Ventricle; A, Atrium; OFT, cardiac outflow tract). Scale bars, F, 50 μm

### Myofibril structure is destabilized in *fstl1b* homozygous mutant ventricular cardiomyocytes

The profound reduction in ventricular contractility prompted us to examine *fstl1b* mutant heart morphology and muscle structure during development. The total number of cardiomyocytes in each chamber was not affected (Fig. 5E,F,G). However, the *fstl1b* homozygous mutant’s ventricular chambers were dysmorphic and exhibited a much smaller size while the atria were dilated (6 out of 9 examined homozygous embryos, Fig. 5A,B) compared to wild type controls (embryos were generated by *fstl1b* heterozygous parents). Normally, by day 3, zebrafish myocardial cells of the ventricle are organized into a single layer of myocardial wall (Fig. 5C, arrowhead). In *fstl1b* mutants, there were multiple layers of cardiomyocytes in the ventricles (Fig. 5D, arrowheads) and the ventricular lumen had collapsed in a more compacted form (comparing Fig. 5A with B and C with D). These data suggested that cell proliferation/division was unlikely to be affected in *fstl1b* homozygous mutant hearts, but that cell structure may be affected.

**Figure 5.**
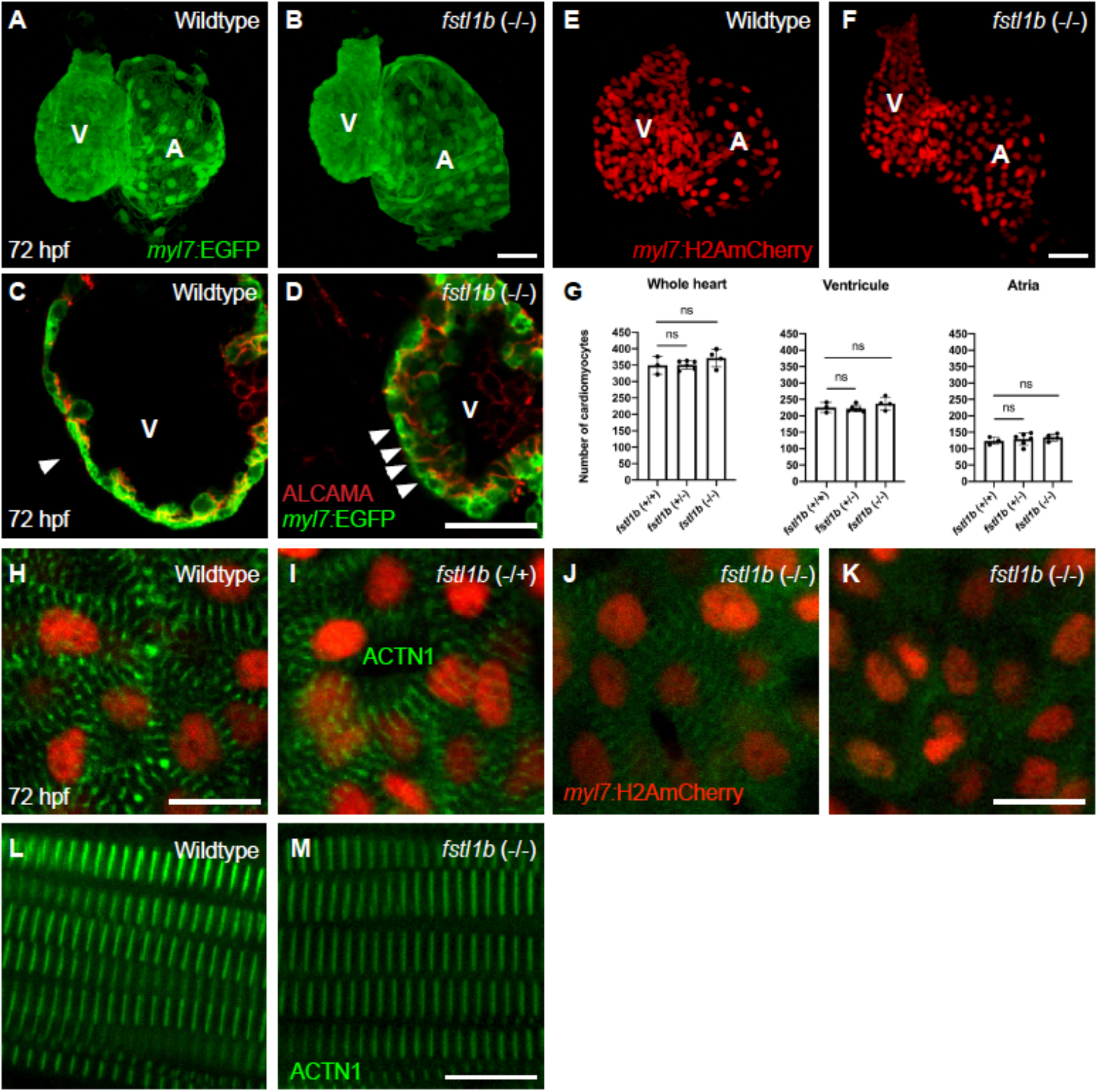
*fstl1b* mutants have a collapsed ventricle during development and exhibit destabilized myofibril structure in cardiomyocytes. (A,B,C,D) Embryonic fish hearts were visualized by EGFP expression in the *myl7:*EGFP transgenic background (green) at 72 hpf, and antibody ALCAMA (ZN5) staining (red in C,D) has been utilized to visualize the cell shape. *fstl1b* homozygous mutant hearts become dysmorphic at three days of development. Especially, the ventricles exhibit a smaller size and seem to be collapsed (B, D), and multiple layers of cardiomyocytes have been observed in the ventricle (D). (V, Ventricle; A, Atrium). (E,F) *myl7:*H2A-mCherry transgenic background has been utilized for counting the cell number of cardiomyocytes during zebrafish development. (G) Comparing to the wild-type controls, *fstl1b* homozygous mutant hearts exhibit a more compacted ventricle, however, there is not significant difference in the cardiomyocyte numbers of both chambers. Significance was determined using one-way ANOVA and Dunnett’s multiple comparisons post hoc test. (ns, not significant). (H,K) Zebrafish hearts stained for alpha-actinin (ACTN1, green) to visualize Z-lines. Comparing to well-defined sarcomere in wildtype controls (H), heterozygous siblings (I), sarcomeres seem to be in disassembled form in *fstl1b* mutant cardiomyocytes with reduced expression of ACTN1 (J) or almost completely absence (I). Although, alpha-actinin is normally expressed in either wildtype (J) or *fstl1b* homozygous (K) skeletal muscle region. Scale bars, A-F, 50 μm; H-M, 10 μm.

To investigate whether Fstl1 signaling activity affects the assembly or the maintenance of sarcomeres in myocardial cells, we examined the distribution of alpha-actinin protein. In striated muscles, alpha-actinin is localized to the Z-line and is a good marker for assessing sarcomere structure. We found that alpha-actinin (ACTN1) expression in the ventricle, detected by immunofluorescence antibody staining, was organized into a banding pattern in both wild type and *fstl1b* heterozygous sibling cardiomyocytes at 72 hpf (Fig. 5H,I). However, half of the *fstl1b* homozygous mutants exhibited reduced expression level of ACTN1 (n=5/10, Fig. 5J) and in some of these homozygotes the ventricular cardiomyocytes with sparse expression had no sarcomeres formed (n=5/10, Fig. 5K). In contrast, the atrial (not shown) and skeletal muscle sarcomere organization seems to be normal in *fstl1b* mutants (Fig. 5J,K). Together, these results indicate a requirement for Fstl1 activity in the maintenance of ventricular myofibril integrity and cardiac chamber morphology.

### Bmp Signaling is upregulated in *fstl1b* mutant hearts

Bmp4 expression patterns exhibit progressive restriction to the AVC (Atrioventricular Canal) junction during cardiac development. Bmp4 is expressed along the entire antero-posterior length of the heart at 24 hpf, but by 48 hpf it is restricted to the AVC and is excluded from the maturing ventricle (33, 34). However, in *fstl1b* homozygous mutants, *bmp4* was ectopically expressed in ventricles at 48 hpf (Fig. 6A,B) and remained ectopically expressed in ventricular myocytes at 3 days postfertilization (not shown). In contrast to *bmp4*, ALCAMA (ZN5) adhesion molecule was relatively normally expressed in the AVC of *fstl1b* mutants (Fig. 5D), similar to wild type control embryos (Figure 5C). Normal ALCAMA expression in mutants suggests that the failure of *bmp4* to become AVC-restricted was not merely due to developmental delay, nor to an overall mispatterning of AV (Atrium-Ventricle) boundaries. Together, these results indicate *fstl1b* homozygous mutants hearts achieve normal AV boundary formation but fail to exclude *bmp4* expression from ventricular myocytes.

**Figure 6.**
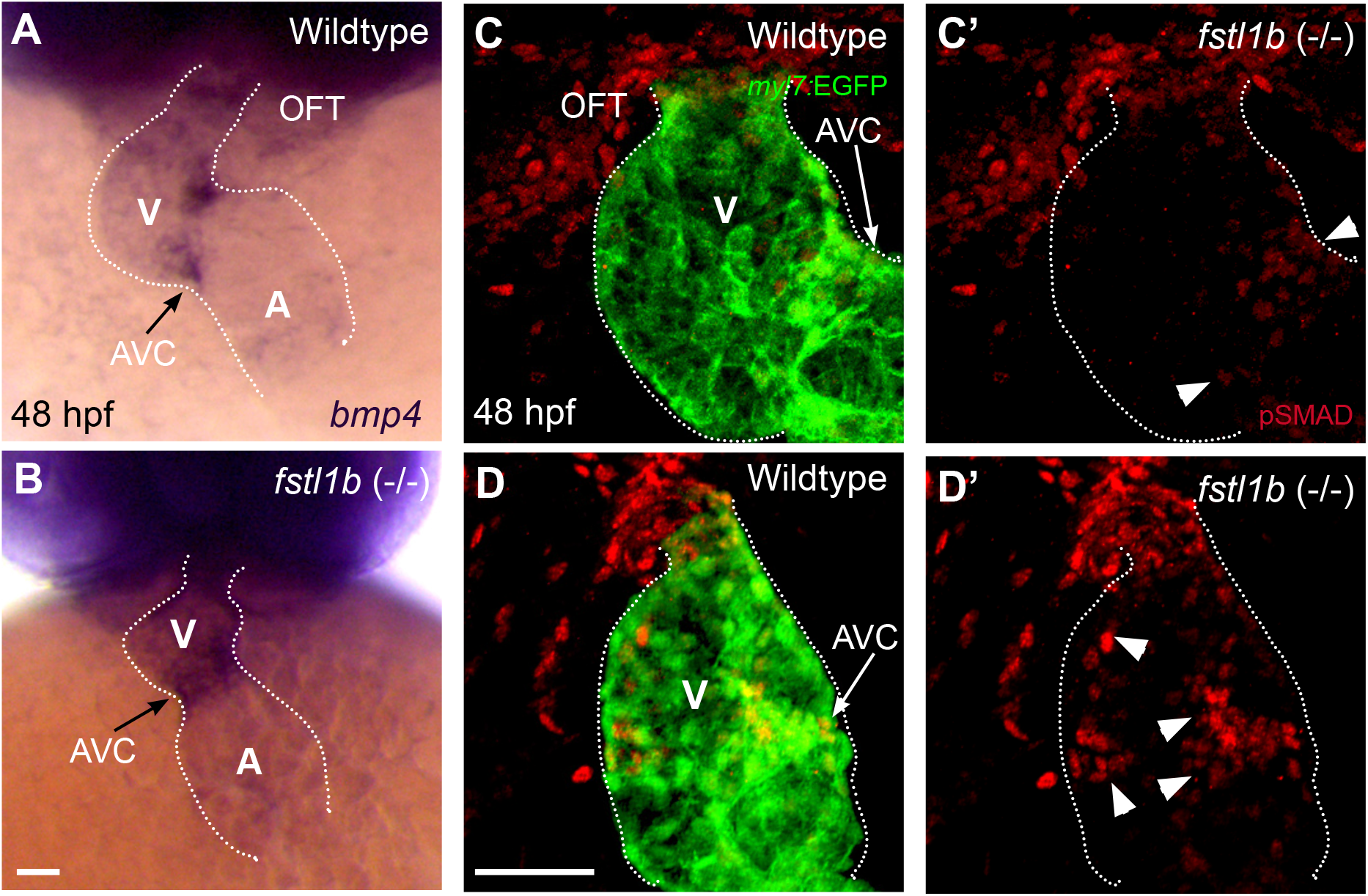
The level of *fstl1b* controls region-specific Bmp signaling in differentiating heart. (A,B) in situ hybridization analysis for *bmp4* expression in the heart at 48 hpf. (A) In wildtype embryos, *bmp4* is primarily restricted in the AVC (Atrium Ventricle Cannel) and OFT (n=4, all exhibited the similar pattern), however *bmp4* expression is less refined at AVC and expanded into the ventricular region in *fstl1b* homozygous mutants (B; n=6, 4 samples exhibited ectopic *bmp4* expression, 2 was normal as seen in wildtype). (C,C’,D,D’) Embryonic fish hearts were visualized by EGFP expression in the *myl7:*EGFP transgenic background (green) at 48 hpf, and antibody P-Smad staining (red) has been utilized to visualize Bmp signaling activities. (C,C’) in wildtype embryos, P-Smad positive cells have been mainly detected in the OFT and AVC (n=5, all exhibited the similar pattern), however P-Smad staining has been observed ectopically in *fstl1b* homozygous mutants’ ventricle (D,D’; n=6, 4 samples exhibited ectopic *bmp4* expression, 2 was normal as seen in wildtype). V, Ventricle; A, Atrium; AVC, Atrium Ventricle Cannal; OFT, cardiac outflow tract. Arrows point to AVC; Arrowhead point to P-Smad positive cells; White dotted lines outline the hearts. Scale bars, 50 μm.

To investigate whether ectopic expression of *bmp4* in the ventricle affects Bmp signaling activity, we performed IHC for phosphorylated-Smad1/5/9 (P-Smad), a downstream marker for Bmp signaling. P-Smad was localized mostly in the nuclei of AVC cells in wild type control hearts (Fig. 6C’,), consistent with *bmp4* transcripts being normally restricted to that region at 48 hpf (Fig. 6A). In contrast, P-Smad was found strongly expressed in the nuclei of ventricular cells in *fstl1b* (-/-) mutant hearts as well as in nuclei of AV canal cells (Fig. 6D’), consistent with the ectopic expression of *bmp4* in the ventricle. These results suggest that the role of *fstl1b* is to confine *bmp4* expression and downstream Bmp response to the AV junction, and to prevent Bmp signaling from spreading into ventricular myocardium at 48 hpf.

## Discussion

In this study we demonstrate that zebrafish *follistatin-like protein 1b* (*fstl1b*) gene has a novel and highly specific function in cardiac development. *fstl1b* homozygous mutants do not show gross defects in early embryonic development, cardiac cell number, and cardiac chamber specification. *fstl1b* is expressed in the developing embryonic heart and required for the normal accumulation/organization of alpha-actinin in the ventricle. The primary physiological effects of *fstl1b* mutations appear to be exerted in the ventricle, including reductions in ventricular contractility and lumen collapse. Alpha actinins belong to the spectrin gene superfamily which represents a diverse group of cytoskeletal proteins, including the alpha and beta spectrins and dystrophins. Alpha actinin is an actin-binding protein with multiple roles in different cell types, and the skeletal, cardiac, and smooth muscle isoforms are localized to the Z-disc and analogous dense bodies, where they forms a lattice-like structure and help anchor the myofibrillar actin filaments and stabilize the muscle contractile apparatus. Previous studies have shown that cardiac contraction can be blocked by increasing Bmp signaling, and can be rescued by reducing the amount of endogenous Bmp4, suggesting that a constrained Bmp signaling in the heart is crucial for normal cardiac contraction (34). The current whole-mount in situ hybridization and immunofluorescence data show that P-Smad, a downstream marker for Bmp signaling, was ectopically induced in *fstl1b* mutant hearts and mimicked the expanded pattern of Bmp signaling in a non-contracting cardiac background (34). Interestingly, in lung development studies, deletion of the mouse *Fstl1* gene resulted in malformed tracheal cartilage and defected alveolar maturation. Significantly, these studies have shown that Fstl1 negatively regulated BMP4/Smad1/5/8 signaling, and is a BMP4 signaling antagonist in controlling mouse lung development (5). Similarly, Fstl1 has been seen as a secreted protein of the Bmp inhibitor class and is required for skeletal organogenesis (4, 6). Altogether, we propose that Fstl1b plays a role in regulating cardiac contractility by constraining *bmp4* expression to the AVC during normal development. The normal expression of *fstl1* is required to stabilize the sarcomeric structure thus ensuring proper contractility. Loss of *fstl1* function results in upregulation of Bmp signaling, which causes the disruption of sarcomere structure and leading to ventricular lumen collapse and non-contraction phenotypes (Fig. 7). Taken together, these observations show that Fstl1 might function as a crucial regulator of Bmp signaling during zebrafish heart development.

**Figure 7.**
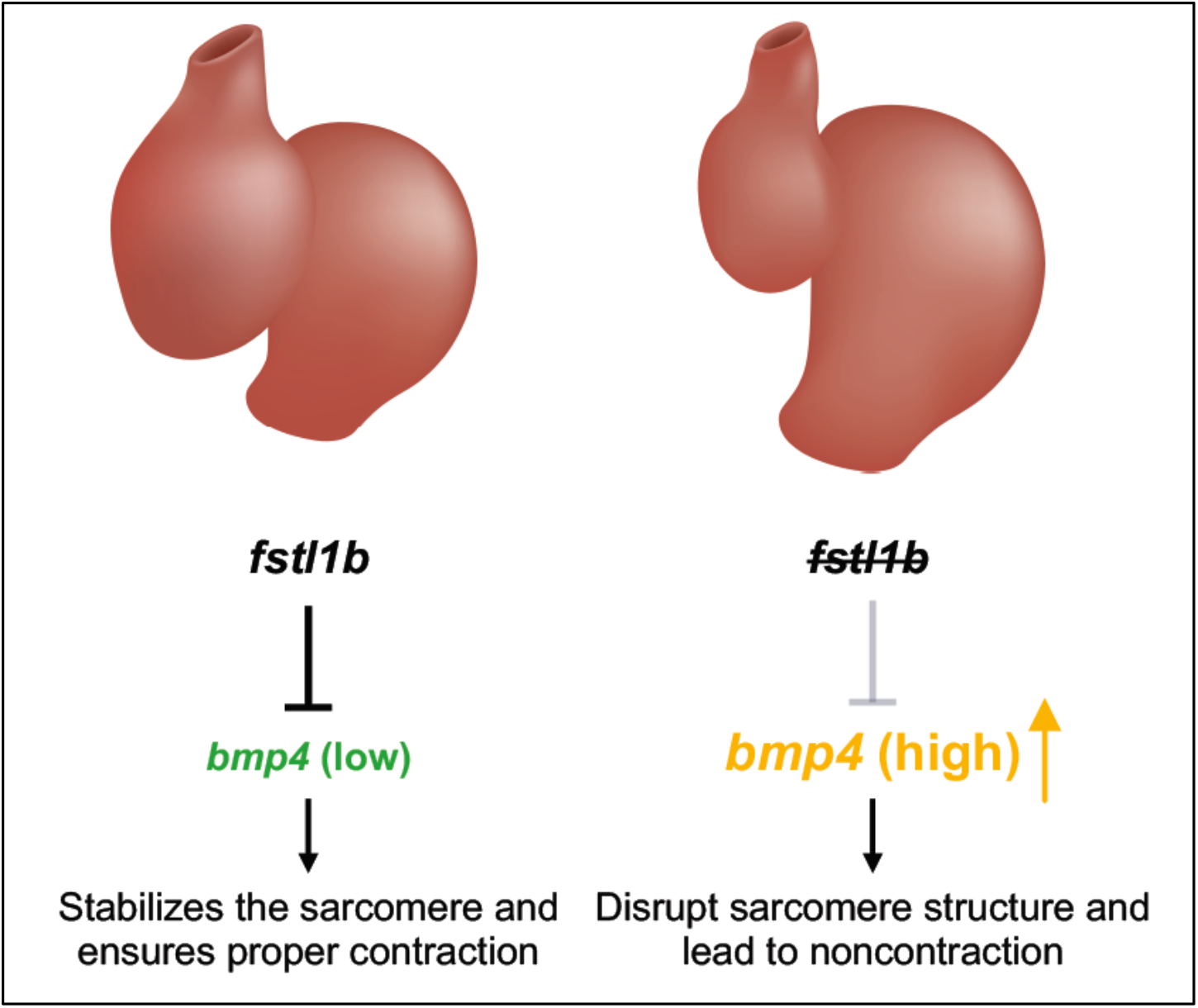
Model for the role of *fstl1b* in cardiac development. Under normal conditions, the normal level of *fstl1b* constrain *bmp4* expression in the ventricle. A relatively low level of *bmp4* stabilizes the sarcomere and ensures proper contraction. Loss of *fstl1* function leads upregulated Bmp signaling, which causes the disruption of sarcomere structure and leads to non-contraction.

It remains unclear how Fstl1 inhibits Bmp signaling in the developing heart. It is known that Fstl1 does not function through a scavenging mechanism, like other known extracellular Bmp inhibitors such as noggin (35). It has been proposed that Fstl1 interferes with BMP receptor complex formation (36, 37), and future challenges will encompass the identification of the factors that determine the mechanisms how Fstl1 prevents the Bmp signaling into the cells during heart development.

## Supporting information

SOHA movie on fstl1b mutant

## Author Contributions

X-X.I.Z. conceptualized project, designed initial experiments, performed experiments, and oversaw all studies.

K.O. and E.E. performed experiments and analyzed the data with X-X.I.Z.

X-X.I.Z., K.O., and E.E. prepared the figures.

K.O. and P.D.S.D. contributed to data interpretation and research funding.

X-X.I.Z. wrote the manuscript with coauthors.

All authors reviewed and contributed to editing the manuscript.

## Acknowledgements

We thank Pilar Ruiz-Lozano and Diane Sepich for excellent comments and suggestions, Nathan Bayardo and Brooke Abrantes for genotyping, Deborah Yelon for *bmp4* antisnse probe plasmid, Jaunian Chen for alpha-actinin whole mount immunostaining protocol, Beth Roman and Arulselvi Anbalagan for technical advice on p-SMAD antibody staining.

## Materials and methods

### Zebrafish

All zebrafish experiments were performed in accordance to protocols approved by IACUC. Zebrafish were maintained under standard laboratory conditions at 28.5 °C. In addition to Oregon AB wild-type, the following transgenic and mutant lines were used: zebrafish carrying the transgenes *Tg(myl7:EGFP)^twu277^* (38), *Tg(myl7:H2A-mCherry)^sd12^* (39).

### SOHA (Semiautomated Optical Heartbeat Analysis)

Larval zebrafish (72 hpf) were immobilized in a small amount of low melt agarose (2.0%) and submerged in conditioned water. Beating hearts were imaged with direct immersion optics and a digital high-speed camera (up to 200 frame/sec, Hamamatsu Orca Flash) to record 30 s movies; images were captured using HC Image (Hamamatsu Corp.). Cardiac function was analyzed from these high-speed movies using semi-automatic optical heartbeat analysis software (30, 40), which quantifies diastolic and systolic intervals, heart period (R-R interval), cardiac rhythmicity, as well as chamber size and fractional area change. All hearts were imaged at room temperature (20-21°C) in a double-blind manner; genotypes were identified after filming and movie analyses had been completed.

Statistical analyses were performed using Prism software (GraphPad). Significance was determined using one-way ANOVA and Dunnett’s multiple comparisons post hoc test.

### Cell counting

To count cardiomyocytes, we used the expression of H2AmCherry in the nuclei (*Tg(myl7:H2A-mCherry*) to qualify as an individual cell, performed the “Spot” function in Imaris to distinguish individual cells in reconstructions of confocal z-stacks (41, 42). Statistical analyses were performed using Prism software (GraphPad). Significance was determined using one-way ANOVA and Dunnett’s multiple comparisons post hoc test.

### Immunofluorescence

Whole-mount immunofluorescence was performed as previously described (15), using primary monoclonal antibodies against sarcomeric myosin heavy chain (MF20). MF20 were obtained from the Developmental Studies Hybridoma Bank maintained by the Department of Biological Sciences, University of Iowa, under contract NO1-HD-2-3144 from NICHD. Anti-sarcomeric alpha-actinin (clone EA53, Sigma-Aldrich, 1:1000), P-Smad1/5/8 (Cell Signaling Technology, 1:100), GFP (anti-Chick, Aves Labs, 1:300), mCherry (anti-Rabbit, Rockland, 1:200), and ZN5 (Developmental Studies Hybridoma Bank, 1:100) were also used for immunostaining. Embryos were classified as having intact sarcomeres if they exhibited at least five adjacent, clearly defined Z-lines marked by a-actinin in any area of the ventricle. Secondary antibody, either donkey anti-chicken AlexaFluor488 (Jackson ImmunoResearch, 1:200) or Donkey anti-Rabbit AlexaFluor568 (Invitrogen, 1:200), was used in 1:200 dilution. Fluorescence images were acquired using an LSM 510 confocal microscope (Zeiss, Germany) with a 40x water objective. Digital images were processed with Adobe Photoshop CS4.

### Cloning of *fstl1a* and *fstl1b* antisense probes

The *fstl1a* and *fstl1b* probe was generated from a plasmid purchased from ATCC (MGC: 101704). The DNA template was amplified using these PCR primers: 5’-TGTACGGAAGTGTTACTTCTGCTC-3’ and 5’-GGATCCATTAACCCTCACTAAAGGGAAGGCCGCGACCTGCAGCTC-3’. The antisense RNA probe was then synthesized with T3 RNA polymerase.

### Standard and fluorescent in situ hybridization

Standard whole-mount in situ hybridization was performed as described previously (43). Fluorescent in situ hybridization was performed using a modified version of published protocols (41, 44). Probes were labeled with digoxigenin and detected by deposition of TSA Plus fluorescein solution (Perkin Elmer). Stained embryos were washed in PBT and mounted in SlowFade Gold anti-fade reagent with DAPI (Molecular Probes) prior to imaging. For fluorescent in situ hybridization in combination with immunofluorescence, we modified a previously described protocol (41, 45). Following probe hybridization, embryos were incubated with both anti-digoxigenin-POD and MF20 antibodies. After detection of digoxigenin-labeled probes by deposition of TSA Plus fluorescein solution (Perkin Elmer), embryos were incubated with an AlexaFluor 647 goat anti-mouse IgG secondary antibody (Invitrogen) to detect MF20 localization.

### Ethics

This study was performed in strict accordance with the recommendations in the Guide for the Care and Use of Laboratory Animals of the National Institutes of Health. All of the animals were handled according to approved institutional animal care and use committee (IACUC) protocols of SBP.

---

**Supplemental Table 1.** A list of available *fstl1a* and *fstl1b* mutant alleles and associated genotyping primer information.

| Gene name | Total exons | sgRNA targeting region | Alleles | Type of mutation | Allele type | Changes in nucleotides | Frame-shifting | Truncated protein | Outcross generation |
| --- | --- | --- | --- | --- | --- | --- | --- | --- | --- |
| *fstl1b* | 11 | Exon 4 | *fstl1b-sbp1* | deletion | predicted null | 4bp | yes | 122/310 aa | F10 |
|  |  |  | *fstl1b-sbp2* | deletion | predicted hypomorphic | 12bp | no | 306/310 aa | F8 |
|  |  |  | *fstl1b-sbp3* | deletion | predicted null | 4bp | yes | 122/310 aa | F8 |
| *fstl1a* | 11 | Exon 2 | *fstl1a-sbp1* | deletion | predicted null | 5bp | yes | 21/314 aa | F8 |
|  |  |  | *fstl1a-sbp2* | insertion | predicted null | 7bp | yes | 18/314 aa | F8 |

Primer sequences information for genotyping:

**Supplemental Figure 1.**
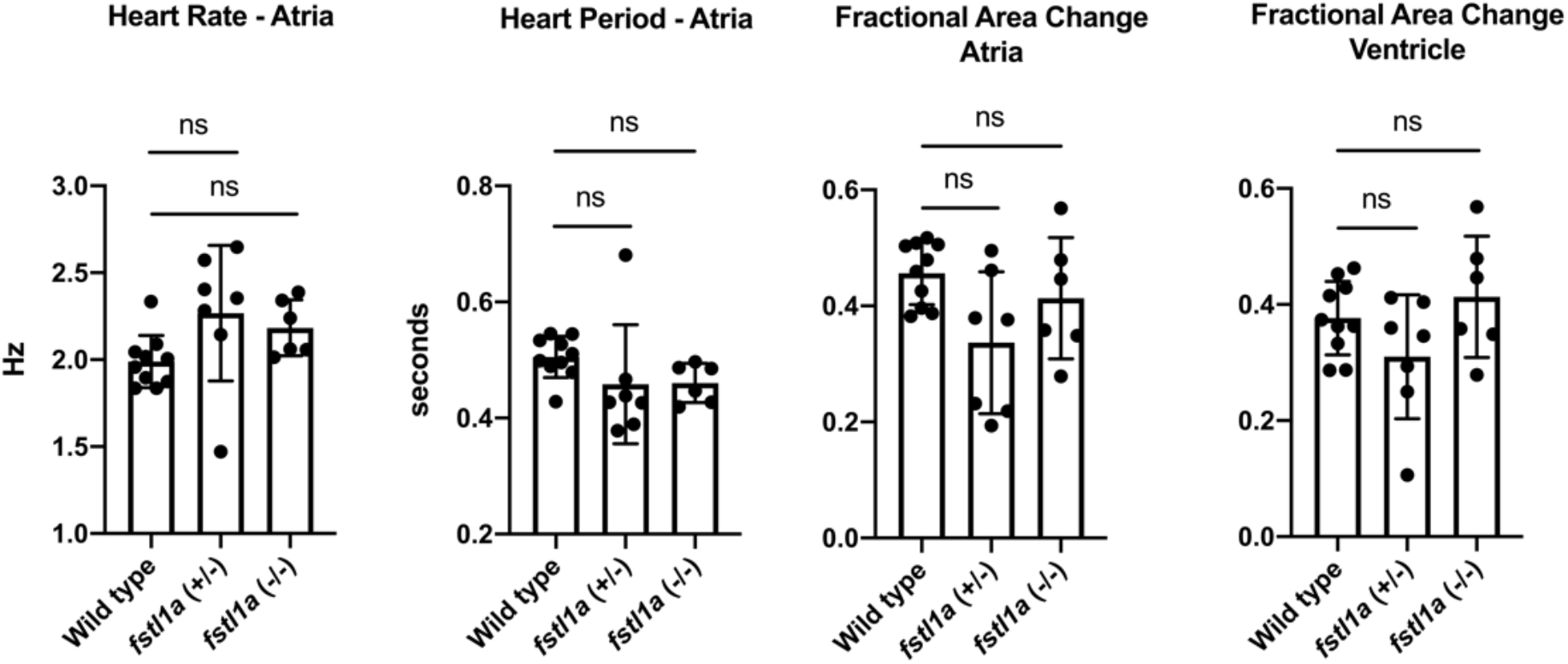
Zebrafish *fstl1a* heterozygous or homozygous mutants exhibit normal cardiac functions. Similar to Figure 3, zebrafish *fstl1a* heterozygous or homozygous mutants’ cardiac function was analyzed from the high-speed movies using semi-automatic optical heartbeat analysis software (SOHA) which quantifies diastolic and systolic intervals, heart period (R-R interval), and fractional area change. All hearts were imaged at room temperature (20-21°C) in a double-blind manner; genotypes were identified after filming and movie analyses had been completed. Statistical analyses were performed using Prism software (GraphPad). Significance was determined using one-way ANOVA and Dunnett’s multiple comparisons post hoc test. (ns, not significant).

## References

1. Shibanuma M, Mashimo J, Mita A, Kuroki T, Nose K. Cloning from a mouse osteoblastic cell line of a set of transforming-growth-factor-beta 1-regulated genes, one of which seems to encode a follistatin-related polypeptide. Eur J Biochem. 1993;217(1):13–9. Epub 1993/10/01. doi: 10.1111/j.1432-1033.1993.tb18212.x. PubMed PMID: 7901004.

2. Zwijsen A, Blockx H, Van Arnhem W, Willems J, Fransen L, Devos K, Raymackers J, Van de Voorde A, Slegers H. Characterization of a rat C6 glioma-secreted follistatin-related protein (FRP). Cloning and sequence of the human homologue. Eur J Biochem. 1994;225(3):937–46. Epub 1994/11/01. doi: 10.1111/j.1432-1033.1994.0937b.x. PubMed PMID: 7957230.

3. Mattiotti A, Prakash S, Barnett P, van den Hoff MJB. Follistatin-like 1 in development and human diseases. Cellular and molecular life sciences: CMLS. 2018;75(13):2339–54. Epub 2018/03/30. doi: 10.1007/s00018-018-2805-0. PubMed PMID: 29594389; PMCID: PMC5986856.

4. Sylva M, Moorman AF, van den Hoff MJ. Follistatin-like 1 in vertebrate development. Birth Defects Res C Embryo Today. 2013;99(1):61–9. Epub 2013/06/01. doi: 10.1002/bdrc.21030. PubMed PMID: 23723173.

5. Geng Y, Dong Y, Yu M, Zhang L, Yan X, Sun J, Qiao L, Geng H, Nakajima M, Furuichi T, Ikegawa S, Gao X, Chen YG, Jiang D, Ning W. Follistatin-like 1 (Fstl1) is a bone morphogenetic protein (BMP) 4 signaling antagonist in controlling mouse lung development. Proceedings of the National Academy of Sciences of the United States of America. 2011;108(17):7058–63. Epub 2011/04/13. doi: 10.1073/pnas.1007293108. PubMed PMID: 21482757; PMCID: PMC3084141.

6. Sylva M, Li VS, Buffing AA, van Es JH, van den Born M, van der Velden S, Gunst Q, Koolstra JH, Moorman AF, Clevers H, van den Hoff MJ. The BMP antagonist follistatin-like 1 is required for skeletal and lung organogenesis. PLoS ONE. 2011;6(8):e22616. Epub 2011/08/10. doi: 10.1371/journal.pone.0022616. PubMed PMID: 21826198; PMCID: PMC3149603.

7. Xu J, Qi X, Gong J, Yu M, Zhang F, Sha H, Gao X. Fstl1 antagonizes BMP signaling and regulates ureter development. PLoS ONE. 2012;7(4):e32554. Epub 2012/04/10. doi: 10.1371/journal.pone.0032554. PubMed PMID: 22485132; PMCID: PMC3317656.

8. Shimano M, Ouchi N, Nakamura K, van Wijk B, Ohashi K, Asaumi Y, Higuchi A, Pimentel DR, Sam F, Murohara T, van den Hoff MJ, Walsh K. Cardiac myocyte follistatin-like 1 functions to attenuate hypertrophy following pressure overload. Proceedings of the National Academy of Sciences of the United States of America. 2011;108(43):E899–906. Epub 2011/10/12. doi: 10.1073/pnas.1108559108. PubMed PMID: 21987816; PMCID: PMC3203781.

9. Ogura Y, Ouchi N, Ohashi K, Shibata R, Kataoka Y, Kambara T, Kito T, Maruyama S, Yuasa D, Matsuo K, Enomoto T, Uemura Y, Miyabe M, Ishii M, Yamamoto T, Shimizu Y, Walsh K, Murohara T. Therapeutic impact of follistatin-like 1 on myocardial ischemic injury in preclinical models. Circulation. 2012;126(14):1728–38. Epub 2012/08/30. doi: 10.1161/CIRCULATIONAHA.112.115089. PubMed PMID: 22929303; PMCID: PMC3548325.

10. Wei K, Serpooshan V, Hurtado C, Diez-Cunado M, Zhao M, Maruyama S, Zhu W, Fajardo G, Noseda M, Nakamura K, Tian X, Liu Q, Wang A, Matsuura Y, Bushway P, Cai W, Savchenko A, Mahmoudi M, Schneider MD, van den Hoff MJ, Butte MJ, Yang PC, Walsh K, Zhou B, Bernstein D, Mercola M, Ruiz-Lozano P. Epicardial FSTL1 reconstitution regenerates the adult mammalian heart. Nature. 2015;525(7570):479–85. Epub 2015/09/17. doi: 10.1038/nature15372. PubMed PMID: 26375005; PMCID: PMC4762253.

11. Chaly Y, Blair HC, Smith SM, Bushnell DS, Marinov AD, Campfield BT, Hirsch R. Follistatin-like protein 1 regulates chondrocyte proliferation and chondrogenic differentiation of mesenchymal stem cells. Ann Rheum Dis. 2015;74(7):1467–73. Epub 2014/03/20. doi: 10.1136/annrheumdis-2013-204822. PubMed PMID: 24641944.

12. Stainier DYR, Fishman MC. The Zebrafish as a Model System to Study Cardiovascular Development. Trends Cardiovas Med. 1994;4(5):207–12. doi: Doi 10.1016/1050-1738(94)90036-1. PubMed PMID: WOS:A1994PK64000002.

13. Gore AV, Monzo K, Cha YR, Pan W, Weinstein BM. Vascular development in the zebrafish. Cold Spring Harb Perspect Med. 2012;2(5):a006684. Epub 2012/05/04. doi: 10.1101/cshperspect.a006684. PubMed PMID: 22553495; PMCID: PMC3331685.

14. Yelon D, Stainier DY. Patterning during organogenesis: genetic analysis of cardiac chamber formation. Semin Cell Dev Biol. 1999;10(1):93–8. Epub 1999/06/04. doi: 10.1006/scdb.1998.0278. PubMed PMID: 10355033.

15. Alexander J, Stainier DY, Yelon D. Screening mosaic F1 females for mutations affecting zebrafish heart induction and patterning. Developmental genetics. 1998;22(3):288–99. Epub 1998/06/11. doi: 10.1002/(SICI)1520-6408(1998)22:3<288::AID-DVG10>3.0.CO;2-2 [pii] 10.1002/(SICI)1520-6408(1998)22:3<288::AID-DVG10>3.0.CO;2-2. PubMed PMID: 9621435.

16. Stainier DY, Fouquet B, Chen JN, Warren KS, Weinstein BM, Meiler SE, Mohideen MA, Neuhauss SC, Solnica-Krezel L, Schier AF, Zwartkruis F, Stemple DL, Malicki J, Driever W, Fishman MC. Mutations affecting the formation and function of the cardiovascular system in the zebrafish embryo. Development. 1996;123:285–92. Epub 1996/12/01. PubMed PMID: 9007248.

17. Dal-Pra S, Furthauer M, Van-Celst J, Thisse B, Thisse C. Noggin1 and Follistatin-like2 function redundantly to Chordin to antagonize BMP activity. Developmental biology. 2006;298(2):514–26. Epub 2006/08/08. doi: 10.1016/j.ydbio.2006.07.002. PubMed PMID: 16890217.

18. Postlethwait JH, Woods IG, Ngo-Hazelett P, Yan YL, Kelly PD, Chu F, Huang H, Hill-Force A, Talbot WS. Zebrafish comparative genomics and the origins of vertebrate chromosomes. Genome Res. 2000;10(12):1890–902. Epub 2000/12/16. PubMed PMID: 11116085.

19. Howe K, Clark MD, Torroja CF, Torrance J, Berthelot C, Muffato M, Collins JE, Humphray S, McLaren K, Matthews L, McLaren S, Sealy I, Caccamo M, Churcher C, Scott C, Barrett JC, Koch R, Rauch GJ, White S, Chow W, Kilian B, Quintais LT, Guerra-Assuncao JA, Zhou Y, Gu Y, Yen J, Vogel JH, Eyre T, Redmond S, Banerjee R, Chi JX, Fu BY, Langley E, Maguire SF, Laird GK, Lloyd D, Kenyon E, Donaldson S, Sehra H, Almeida-King J, Loveland J, Trevanion S, Jones M, Quail M, Willey D, Hunt A, Burton J, Sims S, McLay K, Plumb B, Davis J, Clee C, Oliver K, Clark R, Riddle C, Eliott D, Threadgold G, Harden G, Ware D, Mortimer B, Kerry G, Heath P, Phillimore B, Tracey A, Corby N, Dunn M, Johnson C, Wood J, Clark S, Pelan S, Griffiths G, Smith M, Glithero R, Howden P, Barker N, Stevens C, Harley J, Holt K, Panagiotidis G, Lovell J, Beasley H, Henderson C, Gordon D, Auger K, Wright D, Collins J, Raisen C, Dyer L, Leung K, Robertson L, Ambridge K, Leongamornlert D, McGuire S, Gilderthorp R, Griffiths C, Manthravadi D, Nichol S, Barker G, Whitehead S, Kay M, Brown J, Murnane C, Gray E, Humphries M, Sycamore N, Barker D, Saunders D, Wallis J, Babbage A, Hammond S, Mashreghi-Mohammadi M, Barr L, Martin S, Wray P, Ellington A, Matthews N, Ellwood M, Woodmansey R, Clark G, Cooper J, Tromans A, Grafham D, Skuce C, Pandian R, Andrews R, Harrison E, Kimberley A, Garnett J, Fosker N, Hall R, Garner P, Kelly D, Bird C, Palmer S, Gehring I, Berger A, Dooley CM, Ersan-Urun Z, Eser C, Geiger H, Geisler M, Karotki L, Kirn A, Konantz J, Konantz M, Oberlander M, Rudolph-Geiger S, Teucke M, Osoegawa K, Zhu BL, Rapp A, Widaa S, Langford C, Yang FT, Carter NP, Harrow J, Ning ZM, Herrero J, Searle SMJ, Enright A, Geisler R, Plasterk RHA, Lee C, Westerfield M, de Jong PJ, Zon LI, Postlethwait JH, Nusslein-Volhard C, Hubbard TJP, Roest Crollius H, Rogers J, Stemple DL. The zebrafish reference genome sequence and its relationship to the human genome. Nature. 2013;496(7446):498–503. doi: 10.1038/nature12111. PubMed PMID: WOS:000317984400040.

20. Beumer KJ, Trautman JK, Bozas A, Liu JL, Rutter J, Gall JG, Carroll D. Efficient gene targeting in Drosophila by direct embryo injection with zinc-finger nucleases. Proc Natl Acad Sci U S A. 2008;105(50):19821–6. Epub 2008/12/10. doi: 10.1073/pnas.0810475105. PubMed PMID: 19064913; PMCID: PMC2604940.

21. Tesson L, Usal C, Menoret S, Leung E, Niles BJ, Remy S, Santiago Y, Vincent AI, Meng X, Zhang L, Gregory PD, Anegon I, Cost GJ. Knockout rats generated by embryo microinjection of TALENs. Nat Biotechnol. 2011;29(8):695–6. Epub 2011/08/09. doi: 10.1038/nbt.1940. PubMed PMID: 21822240.

22. Yang H, Wang H, Shivalila CS, Cheng AW, Shi L, Jaenisch R. One-step generation of mice carrying reporter and conditional alleles by CRISPR/Cas-mediated genome engineering. Cell. 2013;154(6):1370–9. Epub 2013/09/03. doi: 10.1016/j.cell.2013.08.022. PubMed PMID: 23992847; PMCID: PMC3961003.

23. Chang N, Sun C, Gao L, Zhu D, Xu X, Zhu X, Xiong JW, Xi JJ. Genome editing with RNA-guided Cas9 nuclease in zebrafish embryos. Cell Res. 2013;23(4):465–72. Epub 2013/03/27. doi: 10.1038/cr.2013.45. PubMed PMID: 23528705; PMCID: PMC3616424.

24. Hwang WY, Fu YF, Reyon D, Maeder ML, Kaini P, Sander JD, Joung JK, Peterson RT, Yeh JRJ. Heritable and Precise Zebrafish Genome Editing Using a CRISPR-Cas System. Plos One. 2013;8(7). doi: ARTN e68708 10.1371/journal.pone.0068708. PubMed PMID: WOS:000321736900106.

25. Hwang WY, Fu YF, Reyon D, Maeder ML, Tsai SQ, Sander JD, Peterson RT, Yeh JRJ, Joung JK. Efficient genome editing in zebrafish using a CRISPR-Cas system. Nature Biotechnology. 2013;31(3):227–9. doi: 10.1038/nbt.2501. PubMed PMID: WOS:000316439500016.

26. Gagnon JA, Valen E, Thyme SB, Huang P, Akhmetova L, Pauli A, Montague TG, Zimmerman S, Richter C, Schier AF. Efficient mutagenesis by Cas9 protein-mediated oligonucleotide insertion and large-scale assessment of single-guide RNAs. PLoS One. 2014;9(5):e98186. Epub 2014/05/31. doi: 10.1371/journal.pone.0098186. PubMed PMID: 24873830; PMCID: PMC4038517.

27. Talbot JC, Amacher SL. A streamlined CRISPR pipeline to reliably generate zebrafish frameshifting alleles. Zebrafish. 2014;11(6):583–5. Epub 2014/12/04. doi: 10.1089/zeb.2014.1047. PubMed PMID: 25470533; PMCID: PMC4248249.

28. Jinek M, Chylinski K, Fonfara I, Hauer M, Doudna JA, Charpentier E. A programmable dual-RNA-guided DNA endonuclease in adaptive bacterial immunity. Science. 2012;337(6096):816–21. Epub 2012/06/30. doi: 10.1126/science.1225829. PubMed PMID: 22745249; PMCID: PMC6286148.

29. Fink M, Callol-Massot C, Chu A, Ruiz-Lozano P, Belmonte JCI, Giles W, Bodmer R, Ocorr K. A new method for detection and quantification of heartbeat parameters in Drosophila, zebrafish, and embryonic mouse hearts. Biotechniques. 2009;46(2):101–+. doi: 10.2144/000113078. PubMed PMID: WOS:000266095700006.

30. Ocorr K, Fink M, Cammarato A, Bernstein S, Bodmer R. Semi-automated Optical Heartbeat Analysis of small hearts. J Vis Exp. 2009(31). Epub 2009/09/18. doi: 10.3791/1435. PubMed PMID: 19759521; PMCID: PMC3150057.

31. Evans SM, Yelon D, Conlon FL, Kirby ML. Myocardial lineage development. Circulation research. 2010;107(12):1428–44. doi: 10.1161/CIRCRESAHA.110.227405. PubMed PMID: 21148449; PMCID: 3073310.

32. Yelon D, Feldman JL, Keegan BR. Genetic regulation of cardiac patterning in zebrafish. Cold Spring Harbor symposia on quantitative biology. 2002;67:19–25. PubMed PMID: 12858519.

33. Beis D, Bartman T, Jin SW, Scott IC, D’Amico LA, Ober EA, Verkade H, Frantsve J, Field HA, Wehman A, Baier H, Tallafuss A, Bally-Cuif L, Chen JN, Stainier DY, Jungblut B. Genetic and cellular analyses of zebrafish atrioventricular cushion and valve development. Development (Cambridge, England). 2005;132(18):4193–204. Epub 2005/08/19. doi: 10.1242/dev.01970. PubMed PMID: 16107477.

34. Samson SC, Ferrer T, Jou CJ, Sachse FB, Shankaran SS, Shaw RM, Chi NC, Tristani-Firouzi M, Yost HJ. 3-OST-7 regulates BMP-dependent cardiac contraction. PLoS Biol. 2013;11(12):e1001727. Epub 2013/12/07. doi: 10.1371/journal.pbio.1001727. PubMed PMID: 24311987; PMCID: PMC3849020.

35. Hirsinger E, Duprez D, Jouve C, Malapert P, Cooke J, Pourquie O. Noggin acts downstream of Wnt and Sonic Hedgehog to antagonize BMP4 in avian somite patterning. Development (Cambridge, England). 1997;124(22):4605–14. Epub 1997/12/31. PubMed PMID: 9409677.

36. Zhou J, Liao M, Hatta T, Tanaka M, Xuan X, Fujisaki K. Identification of a follistatin-related protein from the tick Haemaphysalis longicornis and its effect on tick oviposition. Gene. 2006;372:191–8. Epub 2006/03/07. doi: 10.1016/j.gene.2005.12.020. PubMed PMID: 16517100.

37. Tanaka M, Ozaki S, Osakada F, Mori K, Okubo M, Nakao K. Cloning of follistatin-related protein as a novel autoantigen in systemic rheumatic diseases. Int Immunol. 1998;10(9):1305–14. Epub 1998/10/24. doi: 10.1093/intimm/10.9.1305. PubMed PMID: 9786430.

38. Huang CJ, Tu CT, Hsiao CD, Hsieh FJ, Tsai HJ. Germ-line transmission of a myocardium-specific GFP transgene reveals critical regulatory elements in the cardiac myosin light chain 2 promoter of zebrafish. Dev Dyn. 2003;228(1):30–40. doi: 10.1002/dvdy.10356. PubMed PMID: 12950077.

39. Schumacher JA, Bloomekatz J, Garavito-Aguilar ZV, Yelon D. tal1 Regulates the formation of intercellular junctions and the maintenance of identity in the endocardium. Developmental biology. 2013;383(2):214–26. Epub 2013/10/01. doi: 10.1016/j.ydbio.2013.09.019. PubMed PMID: 24075907; PMCID: PMC3932745.

40. Fink M, Callol-Massot C, Chu A, Ruiz-Lozano P, Izpisua Belmonte JC, Giles W, Bodmer R, Ocorr K. A new method for detection and quantification of heartbeat parameters in Drosophila, zebrafish, and embryonic mouse hearts. Biotechniques. 2009;46(2):101–13. Epub 2009/03/26. doi: 10.2144/000113078. PubMed PMID: 19317655; PMCID: PMC2855226.

41. Zeng XX, Yelon D. Cadm4 restricts the production of cardiac outflow tract progenitor cells. Cell reports. 2014;7(4):951–60. doi: 10.1016/j.celrep.2014.04.013. PubMed PMID: 24813897.

42. Pradhan A, Zeng XI, Sidhwani P, Marques SR, George V, Targoff KL, Chi NC, Yelon D. FGF signaling enforces cardiac chamber identity in the developing ventricle. Development (Cambridge, England). 2017;144(7):1328–38. Epub 2017/02/25. doi: 10.1242/dev.143719. PubMed PMID: 28232600; PMCID: PMC5399623.

43. Thisse C, Thisse B. High-resolution in situ hybridization to whole-mount zebrafish embryos. Nat Protoc. 2008;3(1):59–69. Epub 2008/01/15. doi: 10.1038/nprot.2007.514. PubMed PMID: 18193022.

44. Brend T, Holley SA. Zebrafish whole mount high-resolution double fluorescent in situ hybridization. J Vis Exp. 2009(25). Epub 2009/03/27. doi: 1229 [pii] 10.3791/1229. PubMed PMID: 19322135; PMCID: 2789764.

45. Zhou Y, Cashman TJ, Nevis KR, Obregon P, Carney SA, Liu Y, Gu A, Mosimann C, Sondalle S, Peterson RE, Heideman W, Burns CE, Burns CG. Latent TGF-beta binding protein 3 identifies a second heart field in zebrafish. Nature. 2011;474(7353):645–8. Epub 2011/05/31. doi: nature10094 [pii] 10.1038/nature10094. PubMed PMID: 21623370; PMCID: 3319150.

